# Characterization of the dynamic resting state of a pentameric ligand-gated ion channel by cryo-electron microscopy and simulations

**DOI:** 10.1101/2020.06.19.161356

**Authors:** U Rovšnik, Y Zhuang, L Axelsson, BO Forsberg, V Lim, M Carroni, C Blau, RJ Howard, E Lindahl

## Abstract

Ligand-gated ion channels are critical mediators of electrochemical signal transduction across evolution. Biophysical and pharmacological development in this family relies on high-quality structural data in multiple, subtly distinct functional states. However, structural data remain limited, particularly for the unliganded or resting state. Here we report cryo-electron microscopy structures of the *Gloeobacter violaceus* ligand-gated ion channel (GLIC) under resting and activating conditions (neutral and low pH). Parallel models were built either manually or using recently developed density-guided molecular simulations. The moderate resolution of resting-state reconstructions, particularly in the extracellular domain, was improved under activating conditions, enabling the visualization of residues at key subunit interfaces including loops B, C, F, and M2–M3. Combined with molecular dynamics simulations, the cryo-electron microscopy structures at different pH describe a heterogeneous population of closed channels, with activating conditions condensing the closed-channel energy landscape on a pathway towards gating.

## Introduction

Ligand stimulation of receptors enabling ion flow across cell membranes is a critical mechanism of electrochemical signal transduction in all biological kingdoms. Pentameric ligand-gated ion channels are major mediators of fast synaptic transmission in the mammalian nervous system, but are also found throughout evolution in a variety of biological roles [1]. Members of this protein family share a five-subunit architecture assembled surrounding a central permeation pathway. The extracellular domain (ECD) of each subunit consists of 10 β-strands, with loops A–F enclosing a canonical ligand-binding site [2] at the intersubunit interface. The transmembrane domain (TMD) contains four α-helices (M1–M4), with the M2 helices lining the channel pore, and an intracellular domain of varying length (2–80 residues) inserted between M3 and M4 [3].

Starting from resting (apo-closed) conditions, agonist binding is thought to favor subtle structural transitions to a ‘flip’ or other intermediate state(s) [3], opening of the transmembrane pore, and in most cases refractory desensitization [4]–[7]. Accordingly, a detailed understanding of pentameric channel biophysics and pharmacology requires high-quality structural templates in multiple functional states [8]. Over the last ~12 years several X-ray and cryo-electron microscopy (cryo-EM) structures representing family members from prokaryotes to eukaryotes [3], [9]–[13] have been established. However, few have been solved in the absence of modulatory ligands, leaving substantial unanswered questions as to the structural and dynamic properties of the resting state. The few resting-state structures reported this far have been solved to lower resolution than their liganded counterparts [11], [14], [15], suggesting inherent mobility of this state,which demands models taking the dynamics into account in order to understand its properties and employ it e.g. for drug design.

One of the best studied members of this protein family is the *Gloeobacter violaceus* ligand-gated ion channel (GLIC) [16]–[21]. This proton-gated prokaryotic receptor has proved amenable to crystallization under activating conditions (pH ~4) [22], producing structures in apparent open states in the presence of various ligands, modulators [23], and mutations [24]. Additional structures of GLIC have been reported in lipid-modulated [25] and a so-called “locally closed” state [26], possibly representing gating intermediates. However, only a single GLIC X-ray structure has been reported under resting conditions (pH 7), and only to relatively low resolution (4.4 Å) [27].

Here, we report single-particle cryo-electron microscopy structures of GLIC under multiple pH conditions. To facilitate structure determination and comparison of the same protein in different states, we also employed techniques for density-guided molecular dynamics (MD) simulations [28], validated against manually built models. Although most of the collected cryo-EM data displayed apparently nonconducting pores, comparisons between the reconstructions revealed striking differences in global and local resolution, including local differences in interfacial loops stabilized at different pH. Atomistic simulations of current cryo-EM and previous X-ray structures supported elevated flexibility under resting conditions, particularly in the extracellular domain (ECD). We find increased stability, ECD contraction, and domain twist upon protonation. These results support a broad distribution of conformations under resting conditions, separated by low free energy barriers that produce a dynamic, heterogeneous population of resting-state channels with a closed pore. The distribution narrows upon ligand (e.g. proton) binding to enable transition of a subset of molecules, potentially leading to pore opening.

## Results

### Cryo-EM structures of closed GLIC by manual and automated modeling

To characterize the resting state of the prokaryotic pentameric channel GLIC, we first obtained single-particle cryo-EM data under resting conditions (~pH 7), resulting in a 4.1 Å resolution map (Fig. 1A–B - supplements 1-2, Tab. 1). The map resolution was between 3.5 - 4 Å in the TMD, including complete backbone traces through the C-terminal M4 helix. Side chains in the TMD core were also generally well resolved, including a constriction at the so-called 9’-hydrophobic gate (I233, 2.9 Å Cβ-atom radius) (Fig. 1 supplement 1C), consistent with a closed pore. However, side chains of polar residues E222, N224, and E243 in the pore-lining M2 helix were not as well resolved, and could not be confidently built in the final model. We similarly refrained from building some side chains on the TMD surface, including outward faces of M1 (I198), M3 (M261, E272), and M4 (8 residues), as well as five loop residues between transmembrane helices. Compared to the TMD, the ECD was less well resolved, including some backbone traces that could not be confidently built at the extracellular end (N-terminus and β2–xβ3, β5–β6, and β7–β8 loops), and incomplete side chains distributed throughout the β-strands.

**Figure 1:**
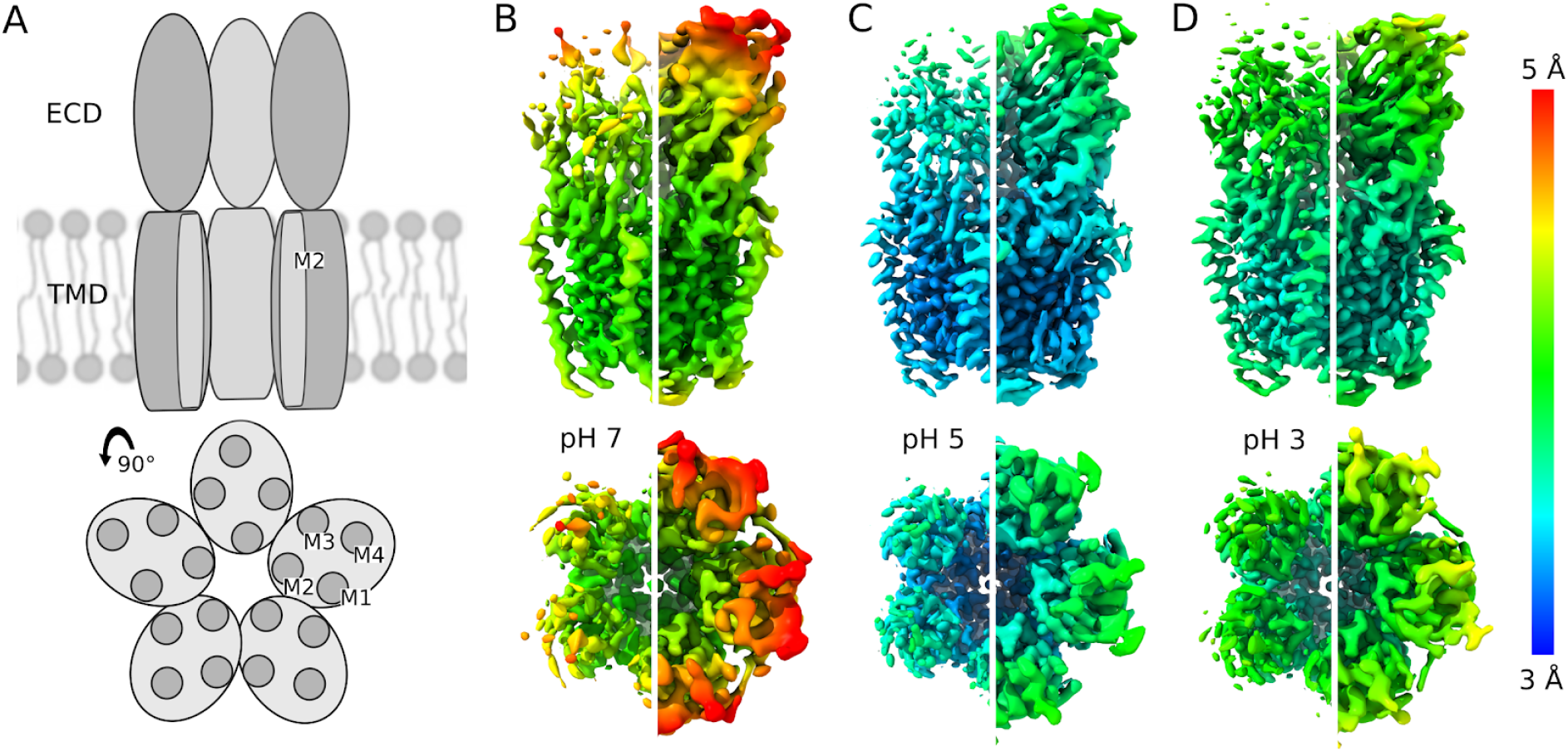
Improvement in local resolution of GLIC cryo-EM densities with decreasing pH. (A) Cartoon representations of GLIC. Upper image is viewed from the membrane plane (side view), with proximal two subunits removed for clarity, revealing the channel axis through the ECD (ovals) and M2 helices within the TMD (cylinders). Lower image shows a 90° rotation viewed from the extracellular side (top view), with transmembrane helices M1–M4 labeled in one subunit. Remaining panels show cryo-EM reconstructions from data collected at (A) pH 7 to 4.1 Å overall resolution, (B) pH 5 to 3.4 Å resolution, or (C) pH 3 to 3.6 Å resolution. Upper and lower images show side and top views respectively, as in panel A. Each density is colored by local resolution according to scale bar at far right, and contoured at both high (left) and low threshold (right) to reveal fine and coarse detail.

**Table 1:**
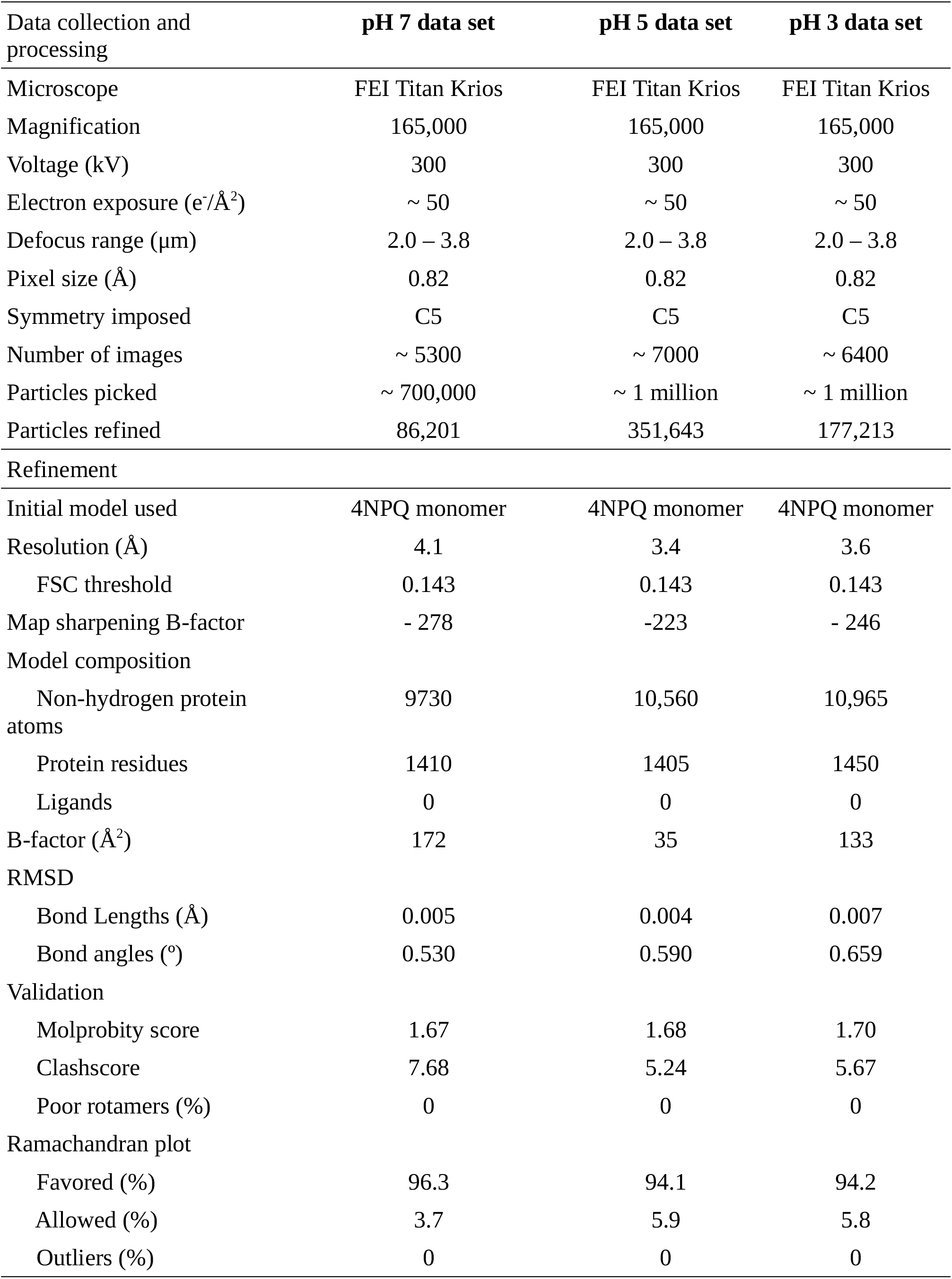
Cryo-EM data collection and model refinement statistics

GLIC has been thoroughly documented as a proton-gated ion channel, conducting currents in response to low extracellular pH with an EC50 around pH 5 [16]. Taking advantage of the capacity of cryo-EM to capture structures under different buffer conditions, we obtained additional reconstructions under partial and maximal (pH 5 and pH 3) activating conditions, producing relatively high-resolution (3.4 Å and 3.6 Å) maps (Fig. 1C-D - supplements 1-2, Tab. 1). To make a more quantitative comparison, a subset of particles was also selected randomly from each dataset collected at pH 5 and 3, containing the same number of particles as the autorefined set from pH 7. Refinements from these low-pH subsets again produced reconstructions to higher resolution than at neutral pH, particularly in the ECD (Fig. 1 - supplement 3), indicating that the varied resolution across pH could not be immediately attributed to particle numbers.

To standardize model building under parallel pH conditions, we also took advantage of new density-guided methods [28], [29] recently implemented in the GROMACS MD package [30], [31]. After equilibrating our pH 7 cryo-EM structure in a lipid bilayer with 150 mM NaCl, we fit this system to masked half-maps obtained at each pH using an automated adaptive-force scaling approach (Fig. 3A). Output models were evaluated relative to the independent half-map at 100 ps intervals, and for all three conditions they converged to stable ensembles within 20 ns (Fig. 3 supplement 1). Final models were obtained by simulated annealing of the best-fit configuration. Because symmetry was not applied during automated fitting, variations could be observed particularly in side chain orientations in different subunits. Given the relative novelty of this automated approach, we also manually built independent models in each condition, which correspond closely (within 2.0 Å Cα root-mean-square deviation; RMSD) to simulation-guided versions (Fig. 3D, Tab. 2).

**Table 2:** Model statistics and validation for models generated with density guided-guided simulations and an RMSD comparison of automatic and manually fitted model.

| Density guided generated models | pH 7 data set | pH 5 data set | pH 3 data set |
| --- | --- | --- | --- |
| RMSD |  |  |  |
| Bond Lengths (Å) | 0.0151 | 0.0184 | 0.0156 |
| Bond angles (°) | 2.10 | 2.40 | 2.05 |
| Validation |  |  |  |
| Molprobity score | 1.07 | 1.10 | 1.14 |
| Clashscore | 0.00 | 0.12 | 0.04 |
| Poor rotamers (%) | 1.59 | 1.59 | 2.06 |
| Ramachandran plot |  |  |  |
| Favored (%) | 93.59 | 94.05 | 94.37 |
| Allowed (%) | 6.15 | 5.18 | 4.66 |
| Outliers (%) | 0.26 | 0.78 | 0.97 |
| Fitted model vs. built model |  |  |  |
| RMSD (Å) | 1.72 | 1.75 | 1.38 |

Regardless of pH, the majority of cryo-EM data reconstructed a constricted hydrophobic gate, likely reflecting predominantly closed-state particles. Still, the relatively high-resolution maps at pH 5 and 3 revealed local rearrangements relative to the maps at neutral pH, including densities that could be fully built for side chains throughout helices M1, M2, M3, and the bulk of M4.

### Local rearrangements associated with protonation

In the ECD, several residues missing backbone density became visible in cryo-EM maps obtained at lower pH, especially in the β2–β3 and β5–β6 loops at the extracellular surface. Furthermore, side chains with poor densities at pH 7 could be confidently built in every β-strand and the β6–β7 (Pro) loop at both pH 5 and 3. At pH 3, additional residues or side chains could be built especially in the β4 strand, β7–β8 (loop B) and β9–β10 (loop C), indicating a progressive increase in local stability with decreasing pH. These three regions form a continuous face of the extracellular β-sandwich, implicated in modulation of GLIC [32] and in agonist binding to other pentameric channels [23], [33]. Indeed, previous X-ray structures included residues in the same three regions that were incomplete in the resting state [27], but resolved under activating conditions, with a relative closure of the cleft between loops B and C [22]. Apparent stabilization at low pH in both cryo-EM and X-ray structures thus supports a coordinated contraction of this region associated with gating.

Notably, backbone densities for three residues in β8–β9 (loop F) were less defined in the pH 5 map compared to the otherwise lower-resolution pH 7 map (Fig. 2B). At pH 3, backbone traces and some side chains could again be built in these regions. Loop F has been extensively implicated in gating of pentameric channels [2] including GLIC [34]; it was recently shown by mutagenesis and covalent labeling to mediate key functional interactions with E35, the main proton sensor in GLIC pH gating [27], which could not itself be built in any of our cryo-EM maps. Thus, successive de- and re-stabilization of loop F under progressively lowered pH was consistent with sequential rearrangements involved in proton sensing.

**Figure 2:**
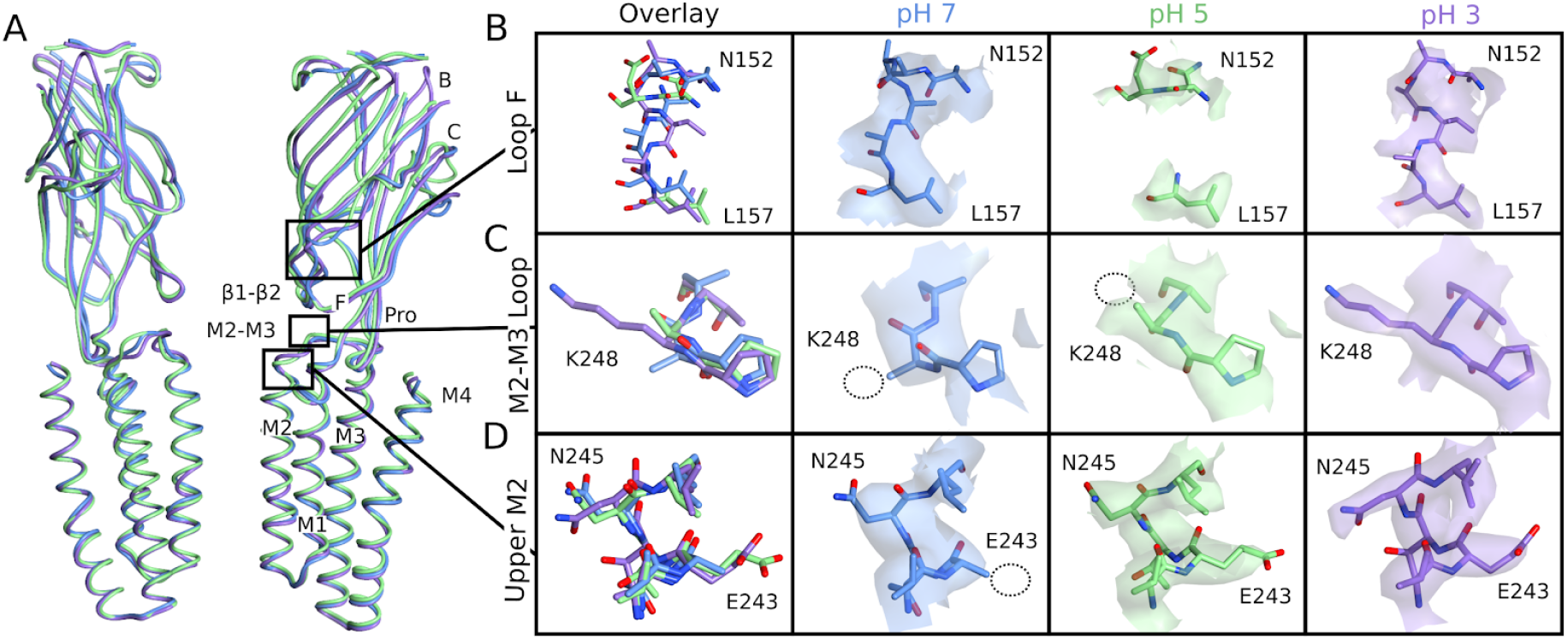
Lower-pH GLIC structures resolve increasing detail. (A) Overlaid side views of backbone ribbons representing two opposing subunits from GLIC cryo-EM models at pH 7 (blue), pH 5 (green), and pH 3 (purple). Functionally relevant features labeled in the righthand subunit include loops B, C, and F and the β1–β2 and Pro loops in the ECD, and helices M1–M4 in the TMD. Remaining panels illustrate local model detail in (B) loop F (N152–L157), (C) the M2–M3 loop (P247–T249), and (D) the upper M2 helix (V242–L246). Leftmost boxes include all-atom overlays at pH 7, 5, and 3, colored by heteroatom with carbons colored as in panel A. Boxes at right show individual models and associated cryo-EM densities. Dotted circles indicate side chains of E243 and K248 that could not be confidently assigned in higher-pH densities; note also that backbone densities were poorly resolved in a portion of loop F (D154–F156) at pH 5.

Although the TMD was generally better resolved under all conditions than the ECD, resolution also improved in low-pH maps for the extracellular mouth of the pore, particularly residue E243 (Fig. 2D, Fig. 3D). This residue has been proposed to stabilize the upper part of the M2 helix during channel opening [27]. In previous X-ray structures, it displayed differential M2-hydrogen bonding patterns at low vs. neutral pH, interacting with T244 of the neighboring subunit under resting conditions [27] but with N245 of the same subunit under activating conditions [22]. Our cryo-EM structures follow a parallel pattern, with E243 contacting T244 at pH 5, but N245 at pH 3.

**Figure 3:**
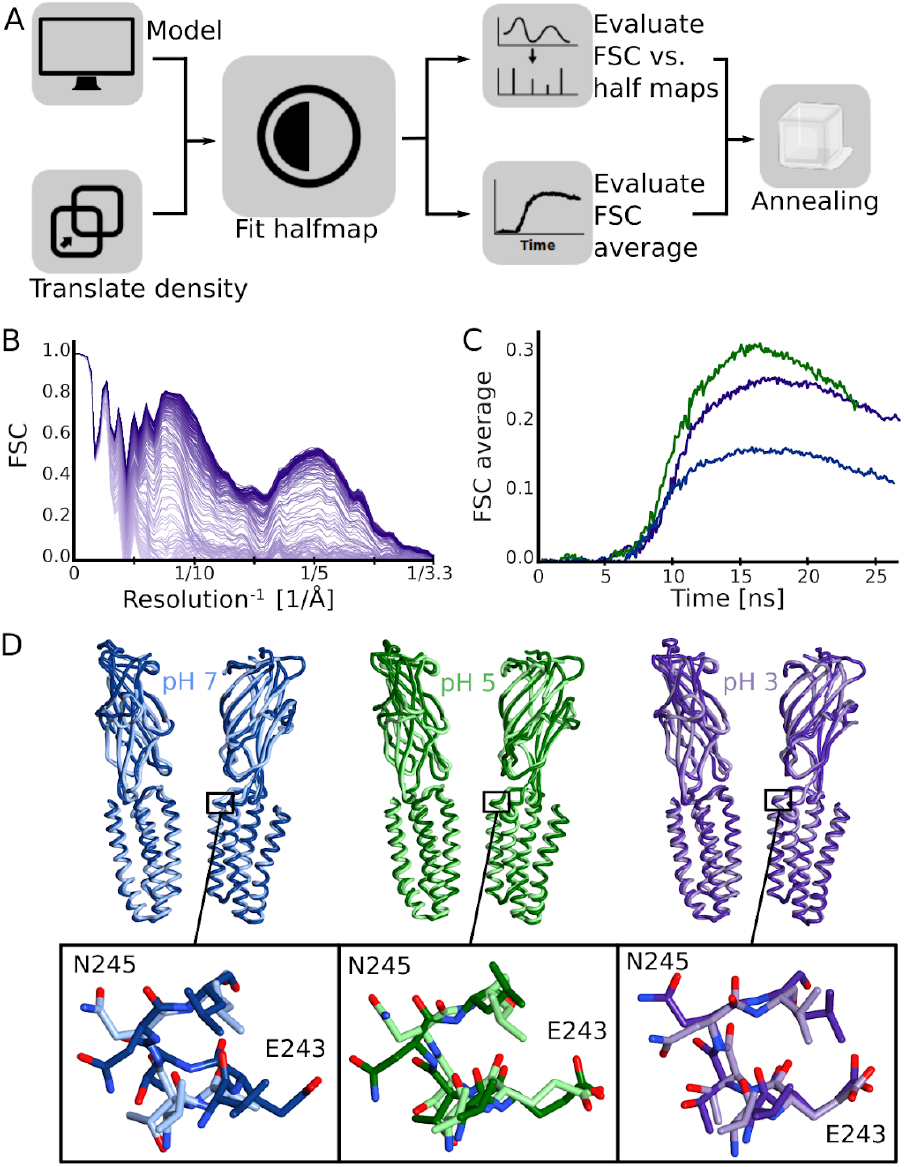
Density-guided fitting enables automated modeling of GLIC structures. (A) Density-guided fitting workflow, starting from an approximate equilibrated model, and ending in a single structure that can be used for comparison and evaluation. (B) Improvement in map-to-model FSC during density-guided fitting to the pH-3 map, showing progress over simulation time (light to dark purple). The FSC varies with resolution, dropping off at the highest overall resolution value for the map (3.4 Å). (C) Average map-to-model FSC over simulation time for fitting to maps at pH 7 (blue), pH 5 (green), and pH 3 (purple). Following steady increases in early fitting stages, each fit reached a maximum similarity around 15 ns, after which the fitting force began to distort the structure. The best-fit frame was chosen for simulated annealing and further analysis. (D) Models from density-guided fitting to data collected at pH 7 (left, blue), pH 5 (center, green), and pH 3 (right, purple). For comparison, manually built models are overlaid in lighter shades of the same color. Upper images show side views of backbone ribbons for two opposing subunits; lower images show detail for the upper M2 helix as in Fig. 2D.

Whereas some residues in the M2–M3 (K248–T249) and M3–M4 (E282–S283) loops remained ill-defined at pH 5 as well as pH 7, all TMD side chains could be built at pH 3. We particularly noted the emergence of side chain density for K248 (Fig. 2D), which was previously reported to orient differently in resting and activating X-ray structures [20], [23]: K248 faced the channel pore and the M2 helix of the same submit at crystallographic pH 7, but reached across the intersubunit cleft to contact N245 on the complementary M2 at pH 4. Similar to low-pH X-ray structures, K248 in the pH 3 cryo-EM structure faced the subunit interface, although the closed conformation of M2 precluded direct contact with N245. In automated density-guided fitting models, K248 was oriented towards the pore in a majority of subunits fit to the pH 7 map, but towards the subunit interface in a majority of subunits fit to pH 5 and 3 data. In several cases, lower-pH conditions produced an upward intersubunit orientation, including electrostatic contacts with complementary ECD residues D32 or Y119. These interactions recall previous reports that ECD-crosslinking of K248 stabilizes a locally closed state [27], possibly representing a gating intermediate.

### Flexibility of the resting-state ECD revealed by molecular dynamics

Given the differential local resolution and conformations observed upon decreasing pH, we asked whether these properties might reflect a flexible resting state, which is systematically stabilized through rearrangements upon protonation particularly in the ECD. To this end, we ran quadruplicate 1–μs all-atom MD simulations on each cryo-EM model embedded in a lipid bilayer. The TMD was consistently stable under all conditions, converging to ~2 Å Cα-RMSD within 250 ns (Fig. 4A). Conversely, for the ECD domain, deviation of Cα atoms was substantially higher in simulations of the pH 7 vs. pH 5 structures (<6 Å RMSD vs. <3 Å RMSD, Fig. 4A, upper row). To assess the impact of presumed protonation at low pH, we also ran simulations of lower-pH structures with a subset of acidic residues protonated according to previously published accessibilities [32]. This adjustment had little effect on the overall stability of the pH 5 structure, but indicated further ECD stabilization at pH 3 (Fig. 4A, lower row). In parallel, simulations of X-ray structures showed ECD destabilization at pH 7 compared to pH 4 (Fig. 4B).

**Figure 4:**
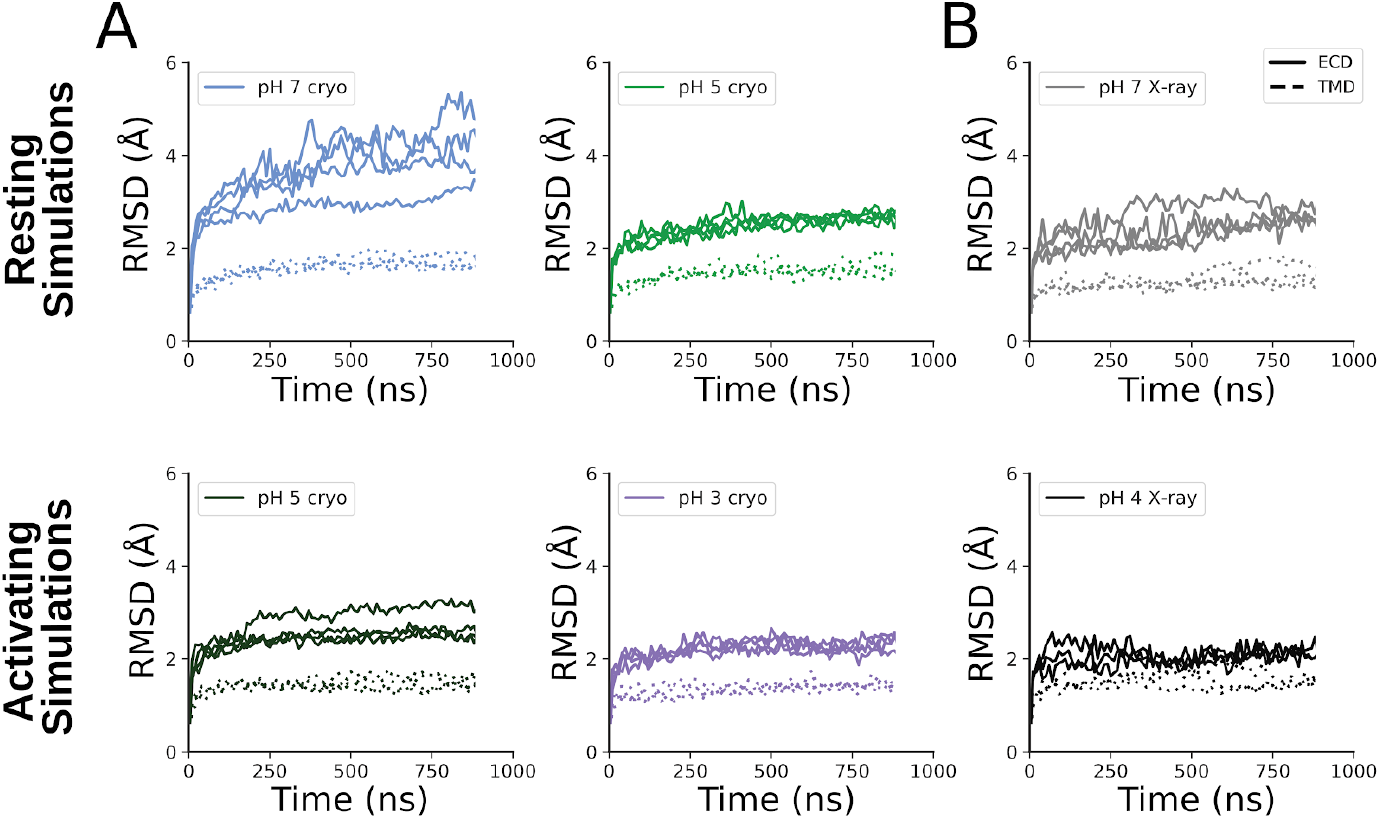
Variable ECD flexibility revealed by MD simulations. (A) RMSD over time for Cα-atoms of the ECD (solid lines) and TMD (dotted lines) in four replicate 1–μs MD simulations of cryo-EM structures determined at pH 7 (blue), pH 5 (light and dark green), and pH 3 (purple). (B) RMSD plots, shown as in panel A, for simulations of X-ray structures at pH 7 (light gray, PDB ID: 4NPQ) and pH 4 (dark gray, PDB ID: 4HFI). Upper images in both panels represent simulations in neutral-pH resting conditions; lower images show simulations in which a subset of acidic residues were protonated to approximate activating conditions.

We further investigated rearrangements in local geometry by monitoring key metrics of gating across MD simulations. First, contraction of the ECD around the channel axis—a reverse “blooming” motion—has been proposed as a key transition on the pathway towards pentameric channel opening [3], [17], [35], [36]. Simulations of the pH 7 cryo-EM structure showed a broad, irregular distribution of the ECD radius, with no clear peak between 30 and 32 Å (Fig. 5A, blue). Conversely, simulations of low-pH cryo-EM structures with either resting or activating protonation featured a more stable and constricted ECD radius, with a single peak around 30 Å (Fig. 5A, green, purple). Previous X-ray structures [22], [27] showed a parallel trend, shifting to a smaller, narrower range in ECD radius values from pH 7 to 4 (Fig. 5A, gray).

**Figure 5:**
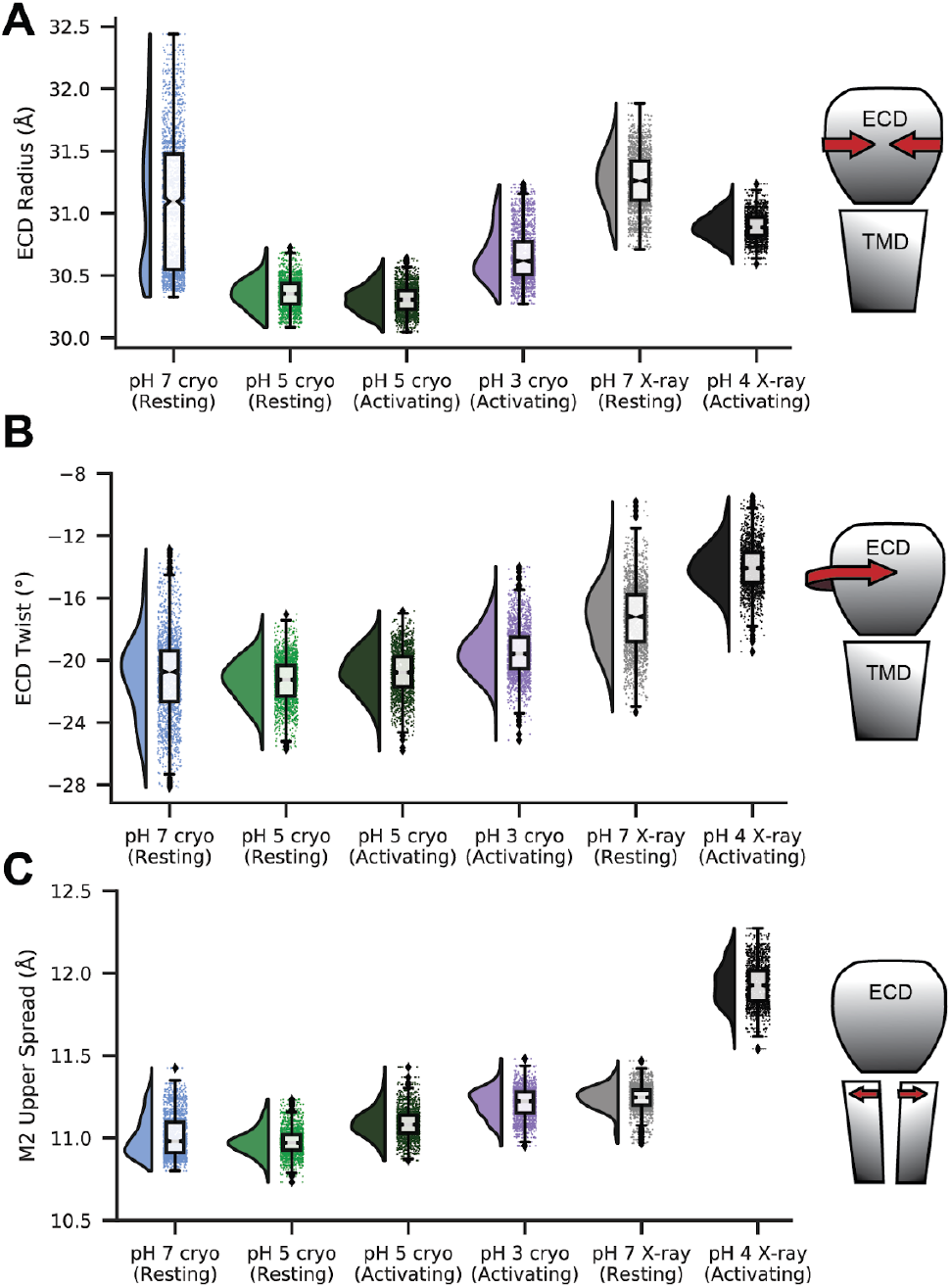
Trends in geometric properties associated with GLIC gating in MD simulations. All panels show raincloud plots [40] for quadruplicate 1–μs MD simulations of cryo-EM structures determined at pH 7 (blue), pH 5 (green) and pH 3 (purple), as well as X-ray structures at pH 7 and 4 (light and dark gray). Simulations were run under deprotonated (resting) or protonated (activating) conditions as indicated, to approximate the experimental state. All plots show probability distributions at left, and raw data plus boxplots indicating sample median, interquartile range, minimum–maximum range, and outliers at right. (A) ECD radius, measured by the average distance from the Cα-atom center-of-mass (COM) of each subunit ECD to that of the full ECD, projected onto a plane perpendicular to the channel axis. (B) ECD twist, measured by the average dihedral angle defined by COM coordinates of 1) a single subunit-ECD, 2) the full ECD, 3) the full TMD, and 4) the same single-subunit TMD. (C) Upper-M2 spread, measured by the average distance from the COM of the upper M2 helix (residues L241–N245) to that of the full TMD.

Similarly, a counter-clockwise twist of the ECD with respect to the TMD has been proposed as an initiating step in pentameric ion channel activation [37], [38], [39]. In parallel to the ECD radius, domain twist fluctuated over a relatively wide range (from −13° to −28°) in simulations of the pH 7 cryo-EM structure (Fig. 5B, blue). This distribution was narrower in simulations of the pH 5 structure, with a single peak around −21° under either resting or activating conditions (Fig. 5B, green). The pH 3 structure exhibited a similarly narrow range, with the peak shifted to around −19° (Fig. 5B, purple). Simulations of previous X-ray structures also showed a relatively broad distribution at pH 7, contracting and shifting to smaller (negative) twist angles at pH 4 (Fig. 5B, gray).

The most prominent geometric change in pentameric channel gating that enables ion flux is the expansion of the transmembrane pore, with the upper region of each M2 helix and the M2–M3 loop spreading outward from the conduction pathway [37]. Simulations of all our cryo-EM structures, as well as the previous pH 7 X-ray structure, featured a relatively contracted upper M2 around 11 Å; in simulations of the pH 4 X-ray structure, upper-M2 spread increased to 12 Å (Fig. 5C). Thus, all major classes obtained by cryo-EM contained similarly closed pores, but exhibited differences in local stability and geometry in the ECD.

## Discussion

The subtle structural changes involved in gating of pentameric ligand-gated ion channels, and their sensitivity to pharmacologically relevant modulation by allosteric drugs, have driven substantial interest in solving structures in multiple key functional states. However, few structures in this family have been reported in unliganded resting states, the presumed “starting block” for gating and modulation. In this context, cryo-EM structures and MD simulations in this work offer insight into the dynamic as well as structural properties of resting and other closed states of GLIC.

Our cryo-EM reconstructions suggest that resting conditions (neutral pH) allow GLIC to sample a rapidly exchanging collection of closed conformations, while activating conditions produce a more discrete population, potentially in equilibrium with open and desensitized states (Fig. 6A). Differential resolutions and orientations were particularly observed in a series of loops spanning the subunit interface (Fig. 6B), including ECD loops B and C on the principal subunit, ECD loop F on the complementary subunit (Fig. 2B), the principal M2–M3 loop (e.g. residue K248; Fig. 2C), and complementary upper M2-helix (e.g. E243; Fig. 2D). Whereas all these regions included incomplete residues at pH 7, backbone traces and most side chains could be built in all these regions at pH 3 (Fig. 6B). Resolution improvements at lower pH, even when controlled for particle number (Fig. 1–figure supplement 3), suggested sampling of a more limited conformational space, including stabilization at the subunit interface.

**Figure 6:**
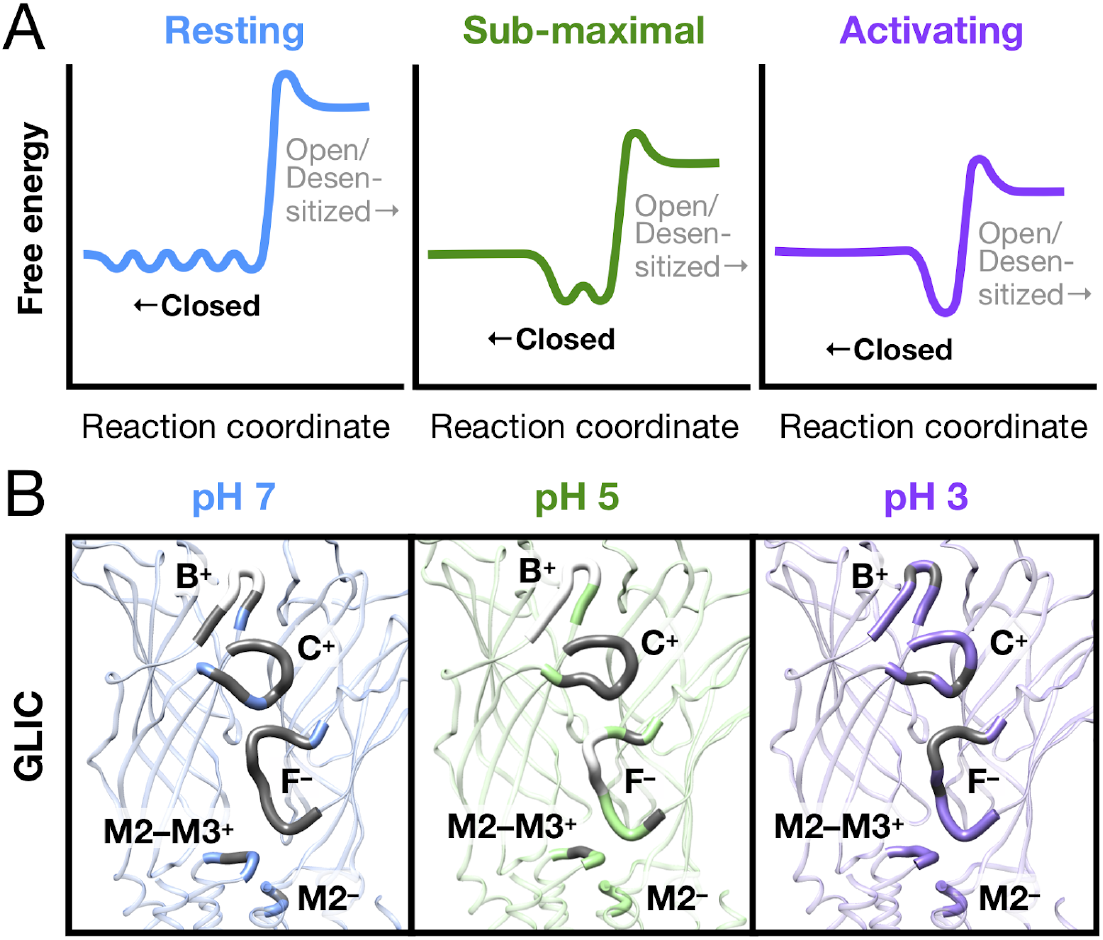
A dynamic resting state of pentameric ligand-gated ion channels. (A) Idealized free-energy diagrams for GLIC, emphasizing closed states. Resting conditions (left, blue) are associated with a rapidly exchanging population of closed channels. Introduction of sub-maximal (center, green) or fully activating conditions (right, purple) condenses the closed free-energy well to a more stable, discrete population, possibly primed for opening. The number of energy wells is meant to represent qualitative progressive consolidation to a discrete population of closed channels upon protonation; details of intermediate, open, and desensitized states to the right of the activation energy barrier are not observed in this work. (B) Regions of the GLIC subunit-interface exhibiting differential resolution in cryo-EM maps reconstructed at pH 7 (left, blue), pH 5 (center, green), or pH 3 (right, purple). Each panel shows the interface between two representative subunits modeled by density-guided MD simulations, viewed from the upper membrane plane. Ribbons for key subunit-interface regions, including loop B, loop C, and the M2–M3 loop of the principal (+) subunit, and loop F and the upper M2 helix of the complementary (–) subunit, are labeled and widened for emphasis. In these regions, residues for which map resolution was insufficient to build complete side chains or backbone atoms are shown in gray and white, respectively.

Transition from a broad, flexible population to a discrete equilibrium upon agonist binding appears to be a consistent pattern in the growing catalog of pentameric channel structures. To our knowledge, wherever such parallel structures have been reported in this family, similarly broad and more flexible structures were observed in resting compared to activating conditions. Within prokaryotes, X-ray structures of the pathogen channel ELIC have been solved to 3.09 Å without ligands [41], but as high as 2.59 Å bound to positive modulators [42]. Similarly, cryo-EM structures of an apo form of ELIC were recently reported in lipid nanodiscs to reach 4.10 Å resolution, while an agonist-bound form was resolved to 3.30 Å in the same work [43]. Although the pore is closed in all reported ELIC structures, a decrease in crystallographic temperature (B-)factors was consistent with extracellular stabilization upon ligand binding, particularly in loops B, C, and F (Fig. 6 - supplement 1A). In the symbiote channel DeCLIC, channel opening was recently shown to involve translocation of a novel N-terminal domain (NTD) to contact the canonical ECD, specifically loops B and C; comparison of DeCLIC structures in apparent open vs. closed states showed a coordinated decrease in B-factors around loops B, C, and F upon NTD binding [44] (Fig. 6 - supplement 1B). Eukaryotic channels have generally required extracellular antibodies or other stabilizing ligands for structure determination in closed as well as open/desensitized states, occluding such comparisons. However, for the full-length mouse serotonin-3 receptor, the relatively low resolution (4.31 Å) of the apo cryo-EM structure [11] was greatly improved by binding the inhibitor granisetron (2.92 Å) to the channel [45] without altering the functional state of the pore, as well as by the agonist serotonin (3.32 Å) with coordinated pore expansion [12]. Interestingly, relatively loose packing of the ECD core has been proposed as a gating strategy specific to eukaryotic members of this channel family [46]; substantiation in eukaryotic receptors of the the dynamic patterns we report here for GLIC may prove a valuable extension of the present work.

Sampling ensembles of conformations in simulations restrained to cryo-EM map data is a potentially valuable approach to better understand the heterogeneity of functional states under different buffer or ligand conditions, provided that the heterogeneity is represented in the reconstruction. In this work, template structures were fitted to cryo-EM reconstructions by conducting an MDsimulation with additional forces to increase the model-to-map similarity. The critical challenge is to balance improved fit against retained structural integrity and sampling of the full ensemble of relevant conformations. We addressed this by adaptively changing the fitting strength according to the change in similarity between model and density during the fitting process. A subsequent analysis of average Fourier shell correlation (FSC) [29] across the fitting trajectory allowed assessment of model fit independent of the driving forces in the simulation, and selection of a model frame in each fitting simulation with minimal structural distortions. The local structure quality was then further improved by annealing. This approach enabled automatic fitting of structures into an experimental density without distorting protein geometry or restraining secondary structure, reflected in the Molprobity [47] (Tab. 2) scores. As an added benefit in this system, structural variability was inherently captured in the five GLIC subunits, which were free to sample conformations independently in density-guided simulations. To assess the flexibility and dynamics of the alternative functional states, we also ran non-restrained MD simulations of the final structures.

The apparent nonconducting conformation of the pore in the GLIC cryo-EM structures under activating conditions was initially surprising, but not inconsistent with previous data. Indeed, at least one previous X-ray structure at pH 4 contained partially occupied closed and open states of the pore, consistent with a substantial population of closed channels even under activating conditions [27]. It is plausible that electrostatic conditions might be modified by interaction with the glow-discharged cryo-EM grid or air-water interface, masking the effect of protonation. Still, we consistently noted subtle shifts in stability and conformation, indicating that local effects of protonation were reflected in the major resolved class. We cannot exclude the possibility of nonconductive channels representing a desensitized state, which has been observed in some GLIC functional assays [48], albeit to a lesser degree than in eukaryotic homologs [16]. However, the cryo-EM structure at pH 3 does not resemble desensitized structures of other channel-family members, but rather hews closely to our other cryo-EM structures. A parsimonious explanation of our low-pH models is that a low open probability produces a predominant population of closed channels, even under maximally active conditions. Although the open probability of GLIC in particular pH conditions is not well established, other channels in this family are known to flicker between conducting and nonconducting states even in the presence of agonist [19], consistent with a large population of closed channels. Indeed, verifiable open states have proved difficult to capture in structural biology, possibly due to their relatively high free energy even under maximally activating conditions. Notably, this rationale is distinct from the apparent challenges to capturing resting structures as reported here, which more likely reflect intrinsic flexibility.

Although resolution and temperature factors can be influenced by multiple experimental and molecular properties aside from flexibility, this qualitative pattern presents a compelling case for a conserved free-energy landscape in this protein family where the resting “state” is rather a relative broad and dynamic ensemble of conformations with the unifying feature of a pore that is neither conducting nor desensitized. Activation of the channel (induced either by pH gating or ligand binding) would then correspond to a transition to a more specific/narrow-distribution state that for many channels would be quite short-lived. This model spotlights a fundamental challenge to capturing a channel’s resting/apo state by traditional structure determination methods, due to the low resolution of any large population of heterogeneous molecules. Closed states captured by the addition of inhibitors, antibodies, or other ligands are often found to be more stable, but may also represent only a subset of the conformational ensemble accessible from the resting state. Structural and dynamic data in this work thus shed light on fundamental properties as well as technical challenges involved in dissecting the ligand-gated channel gating landscape.

## Materials and Methods

### GLIC expression and purification

Expression and purification of GLIC-MBP was adapted from protocols published by Nury and colleagues [32]. Briefly, C43(DE3) *E. coli* transformed with GLIC-MBP in pET-20b were cultured overnight at 37° C. Cells were inoculated 1:50 into 2xYT media with 100 μg/mL ampicillin, grown at 37° C to OD600 = 0.7, induced with 100 μM isopropyl–β-D-1-thiogalactopyranoside, and shaken overnight at 20° C. Membranes were harvested from cell pellets by sonication and ultracentrifugation in buffer A (300 mM NaCl, 20 mM Tris-HCl pH 7.4) supplemented with 1 mg/mL lysozyme, 20 μg/mL DNase I, 5 mM MgCl2, and protease inhibitors, then frozen or immediately solubilized in 2 % n-dodecyl–β-D-maltoside (DDM). Fusion proteins were purified in batch by amylose affinity (NEB), eluting in buffer B (buffer A with 0.02% DDM) with 2–20 mM maltose, then further purified by size exclusion chromatography in buffer B. After overnight thrombin digestion, GLIC was isolated from its fusion partner by size exclusion, and concentrated to 3–5 mg/mL by centrifugation.

### Cryo-EM sample preparation and data acquisition

For freezing, Quantifoil 1.2/1.3 Cu 300 mesh grids (Quantifoil Micro Tools) were glow-discharged in methanol vapor prior to sample application. 3 μl sample was applied to each grid, which was then blotted for 1.5 s and plunge-frozen into liquid ethane using a FEI Vitrobot Mark IV. Micrographs were collected on an FEI Titan Krios 300 kV microscope with a Gatan K2-Summit direct detector camera. Movies were collected at 165,000x magnification, equivalent to a pixel spacing of nominally 0.82Å. A total dose of 40.8 e-/Å^2^ was used to collect 40 frames over 6 sec, using a nominal defocus range covering −2.0 to −3.8 μm.

### Image processing

Motion correction was carried out with MotionCor2 [49]. All subsequent processing was performed through the RELION 3.1 pipeline [50]. Defocus was estimated from the motion corrected micrographs using CtfFind [51]. Following manual picking, initial 2D classification was performed to generate references for autopicking. Particles were extracted after autopicking, binned and aligned to a 15Å density generated from the GLIC crystal structure (PDB ID: 4HFI [22]) by 3D auto-refinement. The acquired alignment parameters were used to identify and remove aberrant particles and noise through multiple rounds of pre-aligned 2D- and 3D-classification. The pruned set of particles was then refined, using the initially obtained reconstruction as reference. Per-particle CTF parameters were estimated from the resulting reconstruction. Global beam-tilt was estimated from the micrographs. Micelle density was eventually subtracted and the final 3D auto-refinement was performed using a soft mask covering the protein, followed by post-processing, utilizing the same mask. Local resolution was estimated using the RELION implementation.

### Model building

Models were built starting from a template using an X-ray structure determined at pH 7 (PDB ID: 4NPQ [27], chain A), fitted to each reconstructed density. PHENIX 1.15-3459 [52] real-space refinement was used to refine this model, imposing the 5-fold symmetry through NCS restraints detected from the reconstructed cryo-EM map. The model was incrementally adjusted in COOT 0.8.9.1 EL [53] and re-refined until conventionally used quality metrics were optimized under a maintained agreement with the reconstruction. Model statistics are summarized in Tab. 1.

### MD simulations

Manually built cryo-electron microscopy structures, as well as previously published X-ray structures (resting, PDB ID 4NPQ [27]; activating, PDB ID 4HFI [22]), were used as starting models for MD simulations. The Amber99sb-ildn force field [54] was used to describe protein interactions. The protein was embedded in a bilayer of 520 Berger [55] POPC molecules. Each system was solvated in a 14 * 14 * 15 nm^3^ box using the TIP3P water model [56], and NaCl was added to bring the system to neutral charge and an ionic strength of 150 mM.

All simulations were performed with GROMACS 2019.3 [30]. Systems were energy-minimized using the steepest descent algorithm, then relaxed for 100ps in the NVT ensemble at 300 K using the velocity rescaling thermostat [57]. Bond lengths were constrained [58], particle mesh Ewald long-range electrostatics used [59], and virtual interact sites for hydrogen atoms enabled a time step of 5 fs. Heavy atoms of the protein were restrained during relaxation, followed by another 45 ns of NPT relaxation at 1 bar using Parrinello-Rahman pressure coupling [60] and gradually releasing the restraints. Finally, the system was relaxed with all unresolvable residues unrestrained for an additional 150 ns. For each equilibrated system, four replicates of 1 μs unrestrained simulations were generated. Analyses were performed using VMD [61], CHAP [62], and MDTraj [63]. ECD radius, ECD twist, and upper-M2 spread were quantified as in previous studies [21], described in Fig. 5.

### Automated density-guided fitting

Automated fitting into cryo-EM densities was performed by density-guided MD simulations using GROMACS 2020.1 [31]. The manually built pH 7 structure, embedded in a bilayer as described above, was used for fitting to all three data sets. Density-guided simulations were performed using one half-map as a bias. To check for overfitting, goodness-of-fit was evaluated using the other half-map. Alignment of the starting model to the density map was assessed and improved iteratively using VMD [61] and the GROMACS editconf functionality, rotating and translating the structure to ensure correct global alignment prior to starting the simulation.

The density-guided simulations were performed at 300K using adaptive force scaling starting at 10 kJ/mol with a feedback time-constant of 40 ps and a Gaussian transform spread-width of the model-generated density of 2 Å. Map similarities were calculated from their inner products after normalization. This normalization leads to higher at-face force constants that however translate to much lower forces and energies.

Evaluation of the overall fit of the model vs. experimental half-map was performed using FSC average values with a threshold of 3.3 Å each 100 ps (equation 2 in [29]). The model configuration at the largest FSC average was chosen as the starting frame for subsequent simulated annealing, which was performed over 2 ns reducing the temperature from 300K to 1K with a density-guided simulation force constant of 10^24^ kJ/mol, corresponding to the force-constant at the largest FSC average. Molprobity scores [47] were calculated every 0.02 ns and the frame with lowest score was used as the final model.

## Acknowledgments

The authors would like to thank the Swedish Cryo-EM National Facility staff, in particular Julian Conrad, José Miguel de la Rosa Trevin and Stefan Fleischmann from Stockholm and Michael Hall from Umeå, for extensive kind assistance with data collection, modeling and supervision.

## Data Availability

Three-dimensional cryo-EM density maps of the pentameric ligand-gated ion channel GLIC in detergent micelles have been deposited in the Electron Microscopy Data Bank under accession numbers EMD-11202 (pH 7), EMD-11208 (pH 5) and EMD-11209 (pH 3), respectively. Each deposition includes the cryo-EM sharpened and unsharpened maps, both half-maps and the mask used for final FSC calculation. Coordinates of all models have been deposited in the Protein Data Bank. The accession numbers for the three GLIC structures are 6ZGD (pH 7), 6ZGJ (pH 5) and 6ZGK (pH 3). Full input data, parameters, settings, commands and trajectory subsets from MD simulations are archived at Zenodo.org under DOI 10.5281/zenodo.3899726.

---

**Figure 1 – supplement 1:**
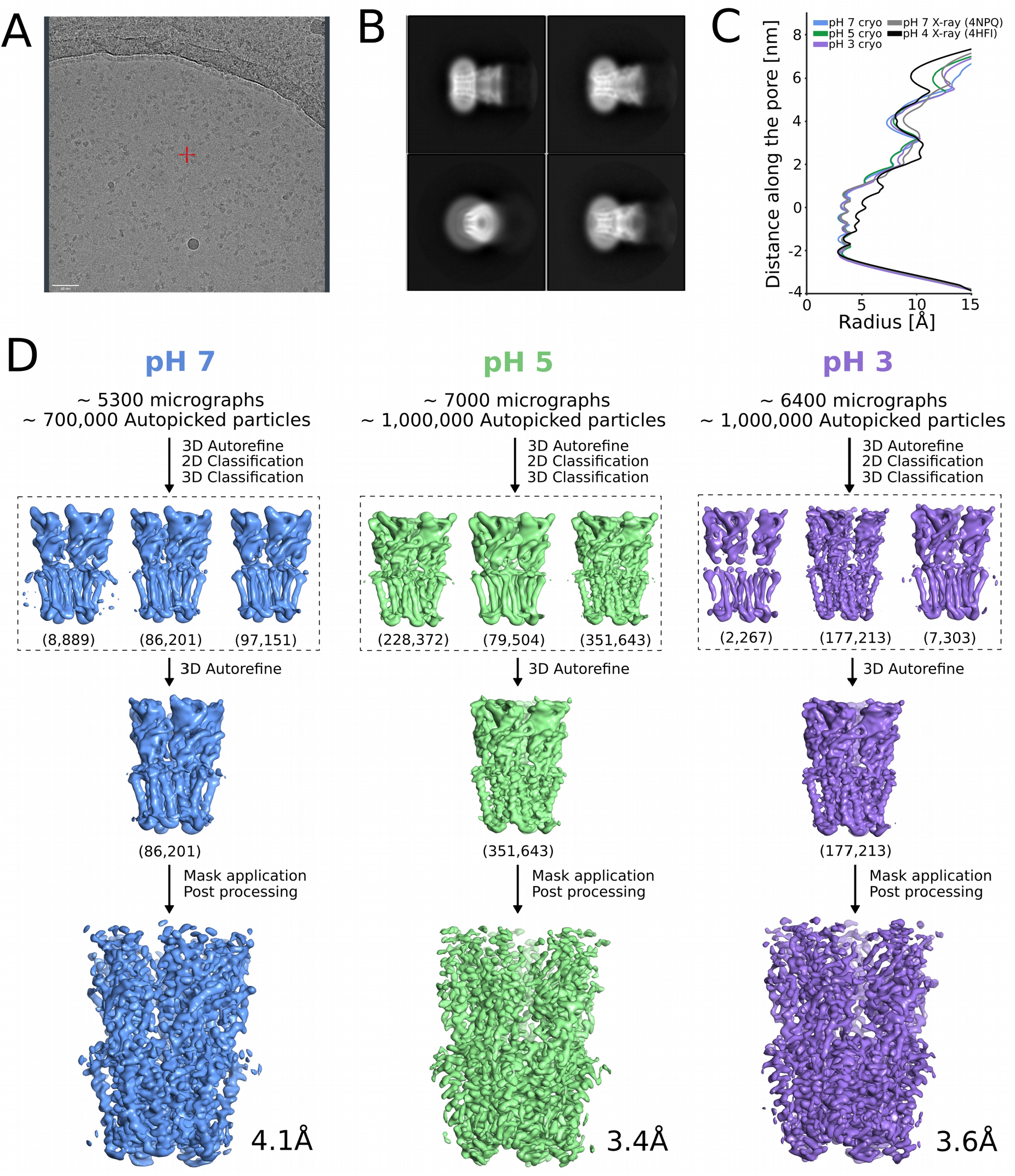
Cryo-EM image-processing pipeline. (A) Representative micrograph from grid screening on a Falcon-3 detector (Talos-Arctica), showing deteigent-solubilized GLIC particles. (B) Representative 2D class averages at 0.82 Å/px in a 256 x 256 pixel box and a 180–Å mask. (C) Radius between Cβ atoms along the GLIC pore for cryo-EM datasets collected at pH 7 (blue), pH 5 (green), and pH 3 (purple), as well as X-ray structure at pH 7 and 4 (light and dark gray). (D) Overview of cryo-EM processing pipelines for data collected at pH 7 (blue), pH 5 (green), and pH 3 (purple) (see Methods).

**Figure 1 – supplement 2:**
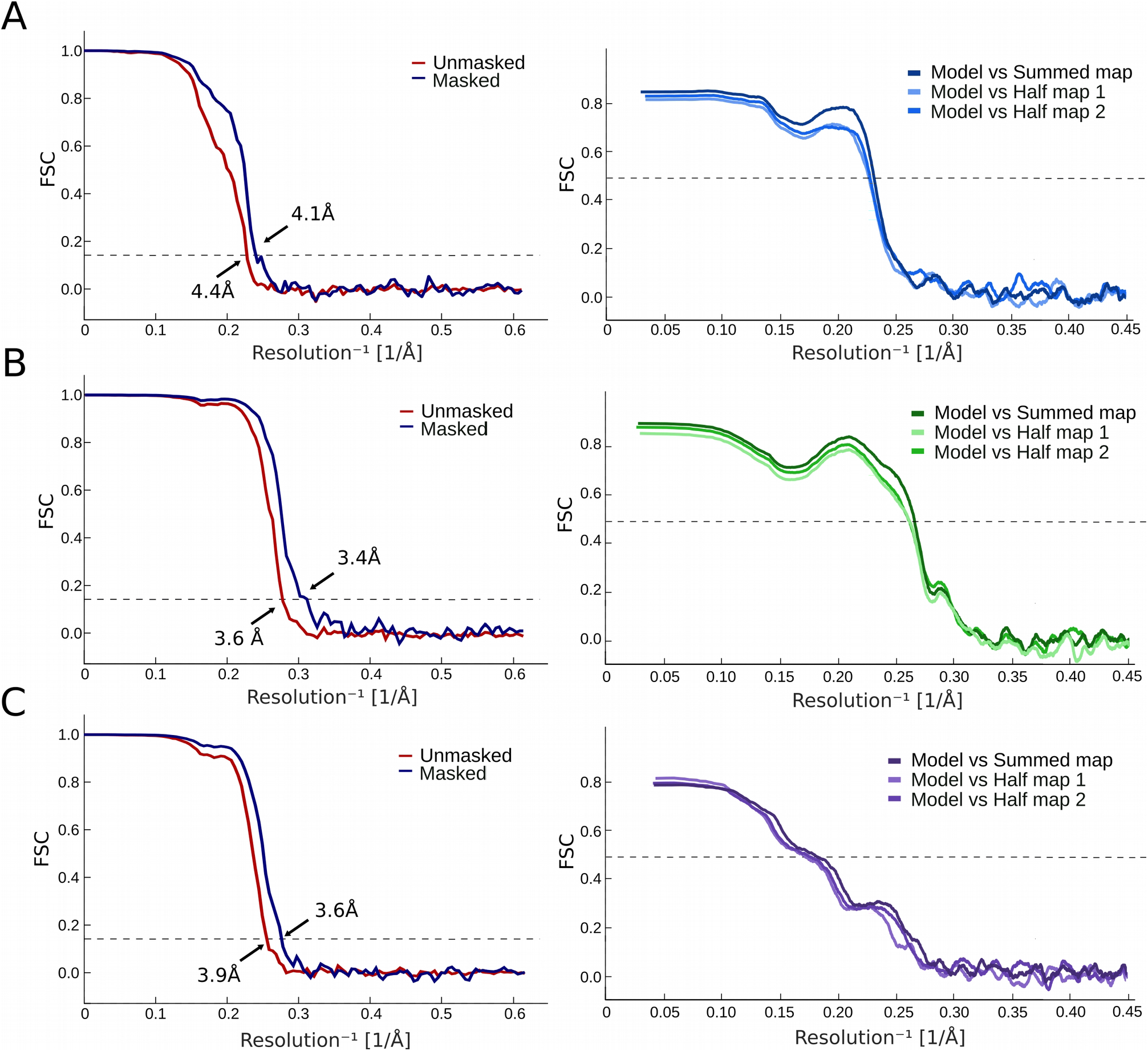
FSC curves and map-to-model FSC for manually modeled cryo-EM reconstructions. *Left:* FSC curves for cryo-EM reconstructions at (A) pH 7, (B) pH 5, and (C) pH 3, before and after applying a soft mask. *Right:* FSC curves for cross-validation between model and half-map 1, model and half-map 2, and model and summed map at (A) pH 7, (B) pH 5, and (C) pH 3.

**Figure 1 – supplement 3:**
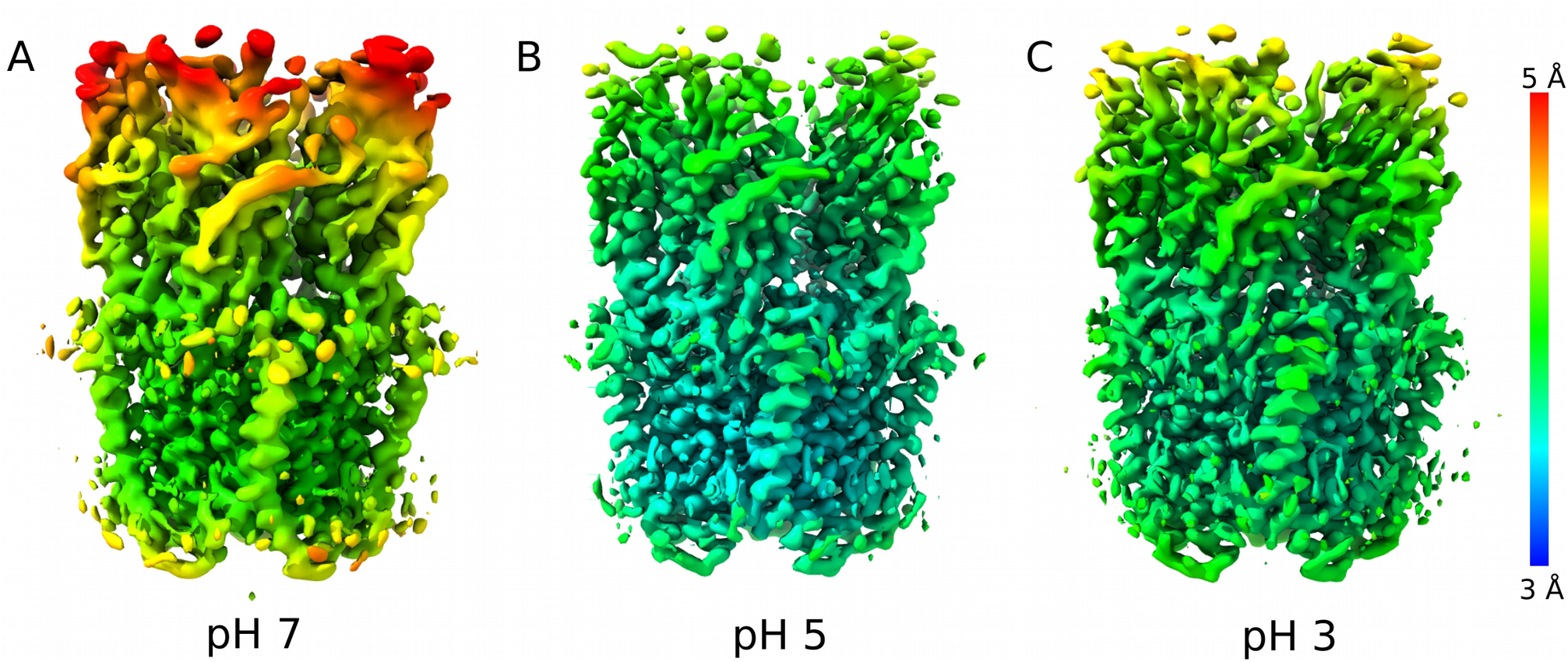
Local resolution of cryo-EM data processed from equivalent particle subsets. Densities at equivalent contour colored by local resolution for reconstructions at (A) pH 7 and data subsets randomly selected to equalize particle number (86,000) at (B) pH 5 and (C) pH 3.

**Figure 3 – supplement 1:**
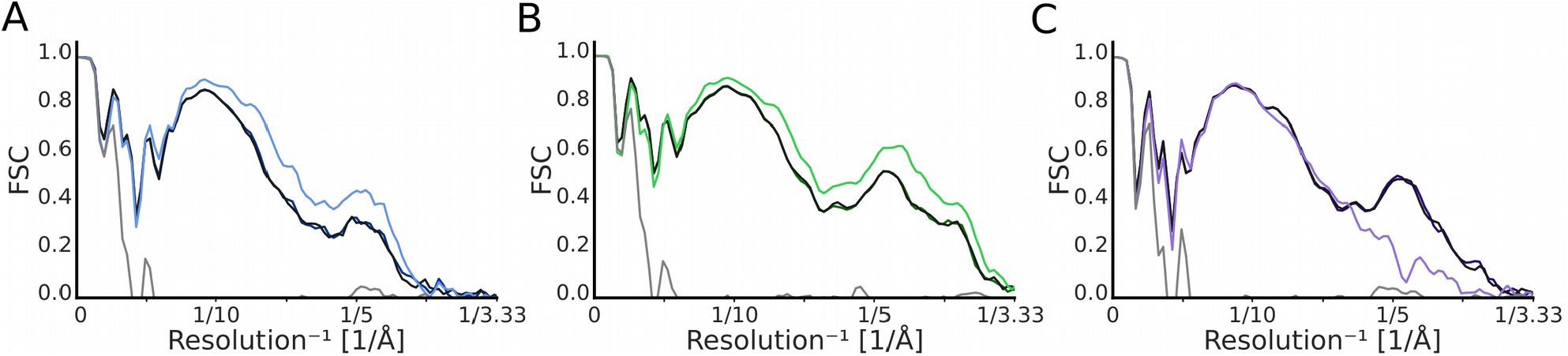
Map-to-model FSC of fitted vs. manual models. Comparison of the map-to-model FSC fit between manually built model and model generated with the automated density-guided simulations. Each of the plots represents an increase of the fit during the course of the fitting process going from light gray to black. The FSC fit for the manually built model for (A) pH 7 is colored light blue and dark blue for the automated model. (B) For pH 5 the manually built model is colored light green and the automated dark green. (C) The FSC fit of the manually built model from the pH 3 dataset is colored light purple and the automated model is colored in dark purple. Note that the FSCs for the fitted are generated from the backbone rather than all atoms, and they are not masked.

**Figure 6 – supplement 1:**
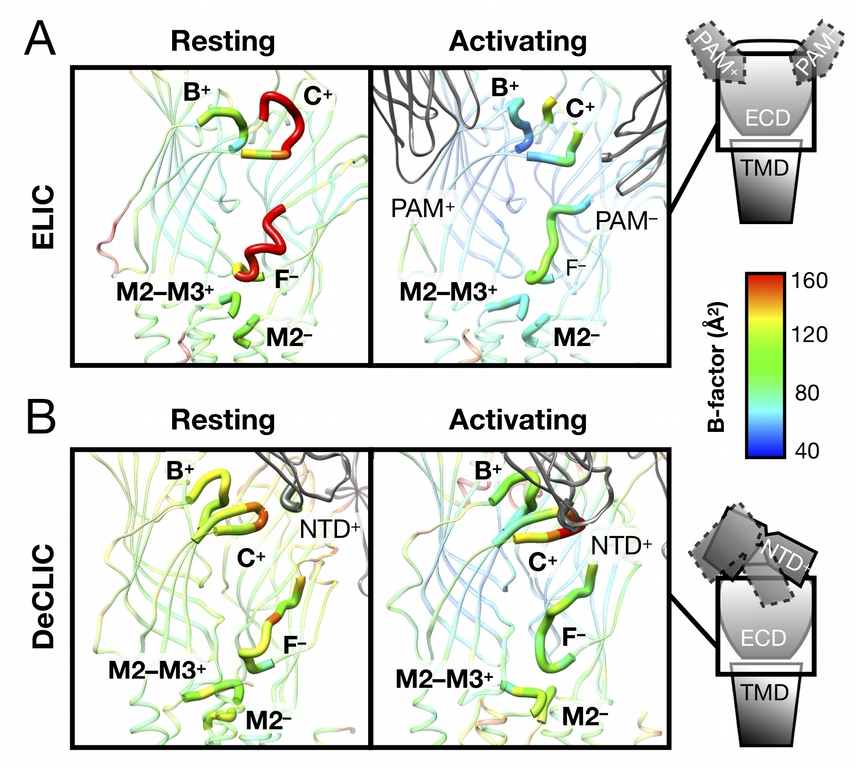
B-factors in homologous channel structures support a conserved resting-state landscape. (A) B-factor distribution in ELIC indicates relative instability prior to ligand binding, particularly in ECD loops B, C, and F. Each panel shows a subunit interface viewed as in Fig. 6B, with residues colored by average B-factor. Comparison of the highest-resolution wild-type X-ray structures reported to date for ELIC in apo (left, PDB ID 3RQU) and liganded states (right, PDB ID 6SSI; complex with modulating nanobody, PAM) indicates lower stability prior to ligand binding, particularly in ECD loops B, C, and F. (B) B-factors in DeCLIC similarly indicate relative flexibility in the ECD under resting conditions. Each panel shows a subunit interface viewed and colored as in panel A. Under non activating conditions (left, PDB ID 6V4S; high calcium), the novel NTD (gray) makes relatively few contacts with ECD loops B and C. In contrast, despite moderately lower overall resolution, the channel under activating conditions (right, PDB ID 6V4S; no calcium) crystallized with generally lower B-factors in the ECD, including interfacial loops.

